# Selective Packaging of Rotavirus Double-Layered Particles into Host Microvesicles

**DOI:** 10.64898/2026.09.16.752160

**Authors:** Zubaida M. Islam, Jonathan Luo, Andy Castillo, Aleksandra Sokol, Emmy Chen, Irin S. Maisha, Ken Nguyen, John J. Dennehy

**Author notes:** Corresponding author: John J. Dennehy.

## Abstract

Rotavirus is a non-enveloped RNA virus traditionally thought to exit host cells through lytic or conventional vesicular pathways, but recent evidence shows that it also exploits host microvesicles for non-lytic transmission. To characterize this transmission pathway, we combined high-resolution microscopy (TEM, confocal microscopy, immunogold labeling) with biochemical and infectivity assays. Our work revealed that rotavirus infection robustly increased microvesicle production, and individual vesicles frequently contained multiple viral particles. Quantitative analysis revealed striking differential partitioning of particle types. Mature triple-layered particles predominated within infected cells, whereas extracellular microvesicles were strongly enriched for immature double-layered particles. Although free double-layered particles lack the outer capsid required for classical receptor-mediated entry, microvesicle-associated particles initiated productive infections, whereas disruption of the vesicular membrane abolished infectivity. These findings suggest an evolutionarily advantageous dual transmission strategy. By using host-derived membranes to transmit otherwise non-infectious intermediates, rotavirus can bypass outer-capsid-dependent entry, potentially reducing the energetic cost of producing fully mature infectious particles while enabling collective transmission. Maintaining a free-virus pathway may nevertheless be important because transmission by individual mature virions imposes population bottlenecks that limit the propagation of defective or cheating genomes and preserve high-fitness genotypes. Thus, partitioning viral progeny between vesicle-associated double-layered particles and free triple-layered particles may balance the immediate benefits of collective spread with long-term genetic quality control. Whether double-layered particle enrichment arises from active sorting or spatial coupling between viral assembly and microvesicle biogenesis remains unresolved.

**Significance:** We demonstrate that rotavirus employs a dual-transmission strategy to optimize viral transmission and replication. While mature triple-layered particles predominate within cells, we show that rotavirus exploits host-derived microvesicles to preferentially package and transmit double-layered particles. This envelopment allows otherwise non-infectious viral intermediates to spread efficiently between cells. Evolutionarily, this reduces the energetic burden of building outer capsids for every genome, freeing resources to maximize viral replication. Concurrently, the virus maintains an independent transmission route via free, mature particles, preventing the accumulation of defective “cheater” genomes inherent to collective vesicle infection. These findings highlight a sophisticated evolutionary balancing act, revealing how viruses deploy structurally distinct particles across varied transmission routes to maximize both local spread and genetic integrity.

## Introduction

Rotavirus (RV), a non-enveloped double-stranded RNA (dsRNA) virus within the *Sedoreoviridae* family, is a leading cause of severe acute gastroenteritis in young children worldwide (1). The infectious virion is structured as a triple-layered particle (TLP) (2). At its center, a VP2 protein shell encapsulates the 11-segment dsRNA genome alongside replication enzymes VP1 (RNA-dependent RNA polymerase) and VP3 (capping enzyme). This inner core is enclosed by a VP6 scaffold to form a transcriptionally active double-layered particle (DLP), which is ultimately capped by an outer layer of VP7 glycoproteins and VP4 attachment spikes to form the mature TLP.

Upon cell entry, the virion sheds its outer capsid, releasing the DLP into the cytoplasm to initiate transcription. Translated viral mRNAs yield the non-structural proteins NSP2 and NSP5, which interact with host lipid droplets and undergo liquid–liquid phase separation to form replication factories called viroplasms (3). Newly assembled inner cores receive their VP6 layer within these factories to become DLPs. To mature into TLPs, DLPs at the periphery of viroplasms interact with NSP4- and VP7-containing COPII-derived membranes that are trafficked from the endoplasmic reticulum and recruited to viroplasms through the autophagy machinery (4). Binding of the cytoplasmic domain of NSP4 to VP6 promotes budding of DLPs through these membranes, producing transiently enveloped intermediates. NSP4 also functions as a viroporin, elevating cytoplasmic Ca²⁺ and promoting autophagy-dependent membrane trafficking during viral morphogenesis (4, 5). VP4 is incorporated during particle maturation, the transient membrane is subsequently lost, and VP7 assembles onto the particle to generate mature, non-enveloped TLPs (5). These virions egress via classical cell lysis or, in polarized cells, through an actin-dependent, Golgi-independent vesicular transport mechanism (6, 7).

However, recent evidence indicates that rotaviruses can also hijack extracellular microvesicles (MVs) for non-lytic egress (8, 9). MVs are 100 to 1,000 nm membrane-bound extracellular vesicles that bud directly outward from a cell’s plasma membrane to transport cargo such as proteins, lipids, and nucleic acids for intercellular communication (10). MV carriage shields the enclosed RVs from host immune defenses and luminal proteases as well as environmental hazards such as chlorine disinfection (8, 11). Importantly, this stealth transport mechanism provides a potential biological mechanism for the frequently observed, yet poorly understood, extra-intestinal dissemination of rotavirus to systemic organs (12–14). Furthermore, packaging multiple virions within a single vesicle benefits the virus by enabling collective infection (8, 11). Allowing multiple viral genomes to simultaneously enter a host cell helps rotaviruses overcome cellular replication barriers and promotes genetic cooperation by facilitating the sharing of replication machinery, which may ultimately drive higher rates of viral reassortment and genetic diversity (8, 15).

Historically, the mature TLP has been considered the sole infectious unit of rotavirus, with the outer capsid proteins VP4 and VP7 deemed strictly necessary for cell entry. In this classical model, the intermediate DLP acts merely as an intracellular transcription engine, incapable of independent transmission. However, during our investigations into MV-mediated egress, we made the unexpected observation that MVs are strongly enriched for DLPs, suggesting the virus utilizes host MVs to transmit an otherwise non-infectious intermediate. If rotavirus actively drives the preferential packaging of DLPs into MVs, it may represent an evolutionary adaptation for efficient host exploitation. We speculate that circumventing the energy-intensive assembly of the outer capsid could allow the virus to redirect the host’s metabolic resources toward genome replication and inner core synthesis. By minimizing the energetic investment required per genome, the virus maximizes the total yield of infectious progeny generated by a single cell. To characterize this non-lytic egress pathway, we used transmission electron microscopy, differential immunogold labeling, and confocal microscopy to structurally quantify the viral payload of rotavirus-induced microvesicles and assess their capacity for receptor-independent infection.

## Results

### Cellular Microvesicle Production Increases Following Virus Infection

To better understand how RV infection impacts cellular MV production, we isolated MVs from rotavirus SA11-infected MA104 cells and uninfected MA104 cells grown in conditioned media using a combination of tangential flow filtration (TFF) and centrifugation at 4, 6, 8, 12, 16, and 24 h post-infection. MV fractions were analyzed for the presence of flotillin-2, an MV marker, using Western blotting (16). Protein concentration was measured using a Qubit 4 instrument (ThermoFisher, Inc.) and normalized before loading. Immunoblot analysis revealed a substantial increase in flotillin-2 protein levels in the infected cells compared to the uninfected controls after 24 h (Fig. 1). Since the flotillin-2 levels in the uninfected controls were too small to be detected even after 24 h, we did a separate Western blot for flotillin-2 levels over 72 h (Fig. 1, inset). These data demonstrate that rotavirus infection robustly induces MV production.

**Figure 1.**
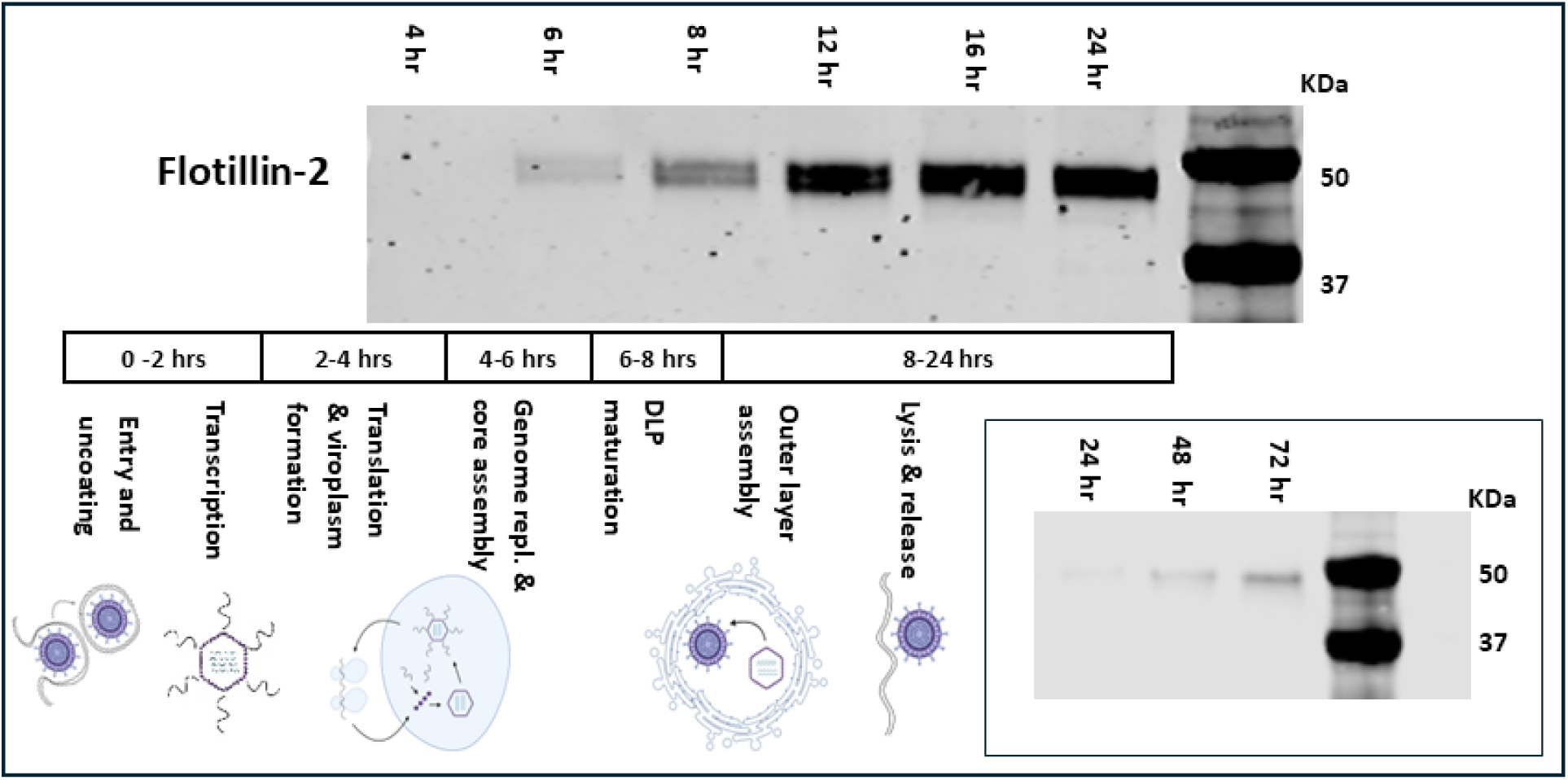
Time-course of virus-induced microvesicle production. MA104 cells were infected with rotavirus strain SA11. MVs were isolated from the culture supernatants at 4, 6, 8, 12, 16, and 24 h post-infection. Sham-infected control cells were prepared and harvested in parallel. Isolated MV fractions were subsequently analyzed by Western blotting using an anti-flotillin-2 primary antibody to quantify relative vesicle yield. The progressive increase in flotillin-2 expression confirms that rotavirus infection upregulates host MV production machinery. Below the Western blot, we show the typical rotavirus life history stage at each timepoint. Due to the absence of a detectable flotillin-2 signal in the sham-infected controls, we repeated the experiment and extended the collection times to 24, 48, and 72 h (Inset).

### Differential Partitioning of Rotavirus Particle Types Between Host Cells and Microvesicles

To quantify the distribution of rotavirus particle types during non-lytic egress, we separated infected cells and extracellular MVs and censused viral particles in both fractions using TEM. Mature rotavirus TLPs possess a smooth outer capsid composed of VP4 and VP7 and have a characteristic diameter of ∼85 nm, whereas immature DLPs lack this outer glycoprotein layer and have a diameter of ∼70 nm (5, 17). The underlying VP6 scaffold is exposed, which produces a rougher, more striated appearance in TEM images (17). We assessed particle diameter and morphology using ImageJ (18). Across 119 micrographs, we characterized 508 intracellular and 664 MV-associated particles and observed a striking segregation of particle types between the two fractions. TLPs predominated intracellularly, accounting for 76.6% of all particles (389 TLPs vs. 119 DLPs). By contrast, DLPs were strongly enriched within MVs and comprised approximately 80% of MV-associated particles (531 DLPs vs. 133 TLPs). A representative TEM image of MV-associated particles is shown in Fig. 2A. The marked enrichment of DLPs within MVs relative to the intracellular population indicates differential partitioning of rotavirus particle types during vesicular release (Fisher’s exact test, p < 0.0001).

**Figure 2.**
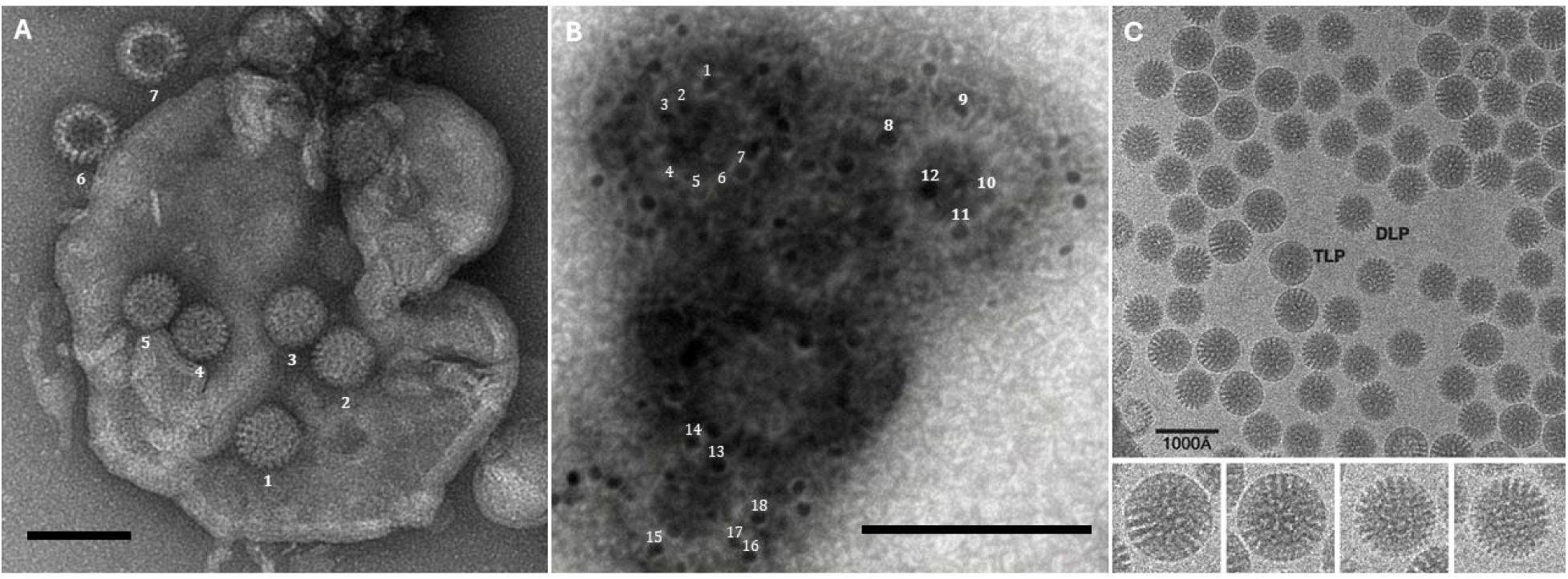
TEM characterization of virus-containing MVs. (A) TEM micrograph showing the morphology and size distribution of SA11 rotavirus particles within a microvesicle isolated from MA104 cells. Total magnification was 20,000x and the scale bar = 200 nm. The measured particle sizes are consistent with double-layered particles (DLPs). Particles 1-7 measured 67, 66, 70, 60, 59, 65, and 76 nm respectively. Note that particles 6 and 7 are likely externally associated with the MV and show evidence of stain intrusion into the capsid due to being DLPs (47). (B) Dual-immunogold labeling of MV fractions. The ∼8-nm gold particles correspond to secondary antibodies recognizing anti-VP6. The ∼8-nm gold particles correspond to secondary antibodies recognizing anti-VP7. Numbers directly above particles correspond to particle size measurements. Particles 1-18 measured 5.6, 5.2, 5.9, 5.8, 7.4, 5.3, 7.1, 8.2, 7.9, 7.5, 8.2, 8.8, 7.0, 7.6, 8.2, 7.3, 7.0, and 7.5 nm, respectively. The particle in the upper left is consistent with a TLP and the particles in the upper right and lower are consistent with DLPs. Total magnification was 30,000x and the scale bar = 100 nm. (C) Cryo-EM micrograph of rotavirus particles showing morphological and size differences between TLPs and DLPs. The larger TLPs exhibit a characteristic smooth, circular profile, whereas the smaller DLPs display a bristly outer surface. Magnified views of two TLPs (left) and two DLPs (right) are provided in the bottom insets. Scale bar = 1000 Å. Adapted from Figure 43 in (48).

### Immunogold Electron Microscopy Identifies MV-Associated Viruses as DLPs

While our initial morphological analysis based on particle diameter strongly suggested the presence of DLPs, we sought definitive molecular confirmation of the virion assembly state within MVs. To achieve this, dual immunogold labeling was performed using primary antibodies specifically targeting either the VP6 intermediate scaffold or the VP7 outer capsid glycoprotein. This differential labeling strategy leverages the highly ordered structural topology of the rotavirus virion. In fully mature TLPs, the dense outer layer of VP7 trimers completely envelops the particle, shielding the underlying VP6 layer from antibody recognition. Consequently, anti-VP6 antibodies will exclusively bind to DLPs where the VP6 proteins remain exposed, whereas anti-VP7 antibodies selectively flag the intact outer capsid of mature TLPs.

To concurrently visualize and differentiate these two distinct populations within the same sample, we utilized secondary antibodies conjugated to colloidal gold nanoparticles of disparate sizes (e.g., ∼ 8 nm gold for anti-VP6 and ∼ 6 nm gold for anti-VP7). TEM analysis of the labeled microvesicles provided clear validation of our prior sizing data. The viral populations sequestered within the microvesicles exhibited dense, localized clusters of the VP6-associated gold label, alongside a conspicuous paucity of the VP7-associated label. Ultimately, this specific labeling profile confirmed a significant enrichment of DLPs over TLPs within the vesicles, strongly supporting the model that EVs are predominantly loaded with DLPs (Fig. 2B).

### Enrichment of VP6 Protein Associated with Double-Layered Particles in Budding Microvesicles

Following the structural characterization of MV-associated viral particles, we investigated the biogenesis of these vesicles in fixed rotavirus SA11-infected MA104 cells. To track the spatial distribution of viral components, we performed confocal microscopy utilizing primary antibodies targeting specific structural proteins: an anti-VP6 antibody to detect the intermediate double-layered particle (DLP) scaffold, and an anti-VP7 antibody to detect the mature triple-layered particle (TLP) outer capsid glycoprotein. These targets were visualized using differentially colored fluorescent secondary antibodies, Alexa Fluor 488 for VP6 and Alexa Fluor 647 for VP7. Confocal imaging revealed a distinct, robust accumulation of the VP6 protein at the plasma membrane, strongly co-localizing with actively budding microvesicles (Fig. 3). It is important to note that while this fluorescent signal confirms the tight association of the VP6 structural protein with these egressing vesicles, the resolution limits of standard confocal microscopy preclude us from definitively concluding whether these accumulations represent fully assembled, intact DLPs or solely the concentrated VP6 protein at these specific membrane sites (Fig. 3).

**Figure 3.**
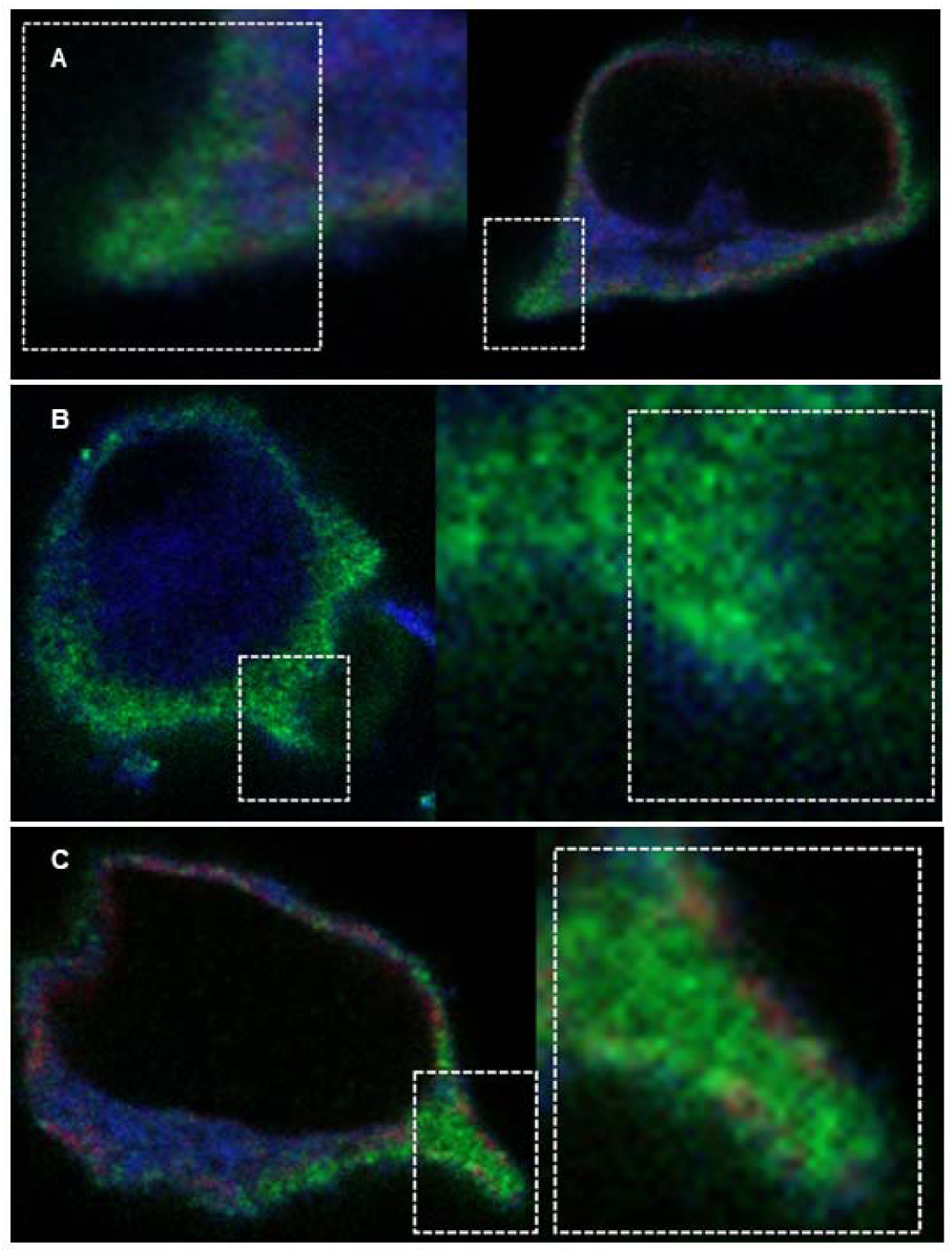
Confocal microscopy of viral budding dynamics. Dual-color immunofluorescence of MV-associated particles at 8 h (Panel A), 14 h (Panel B), and 18 h (Panel C) post-infection. Dashed line boxes indicate areas that were magnified for closer inspection. Double-layered particles (DLPs) were labeled with anti-VP6 (Alexa Fluor 488, green) and triple-layered particles (TLPs) with anti-VP7 (Alexa Fluor 647, red). Peripheral budding sites show a significant accumulation of the DLP signal relative to the TLP signal.

### Double-Layered Particles Recovered from MVs Are Infectious

Having established that the vesicular cargo predominantly consists of DLPs, which are classically considered non-infectious in the extracellular space due to the absence of the VP4/VP7 outer capsid, we next investigated whether these MVs are biologically functional and capable of initiating new infections. To test this, we performed plaque assays on uninfected host cell monolayers using purified MVs. A neutralizing anti-VP7 antibody was used to inactivate residual TLPs (19). The intact virus-laden MVs readily produced distinct viral plaques in the absence of trypsin, a protease typically required to activate free rotavirus virions by cleaving the VP4 attachment spike (Fig. 4 A and B)(9, 20–22). This experiment demonstrates that the observed infectivity is strictly mediated by the vesicular envelope rather than the traditional, receptor-dependent entry pathway.

**Figure 4.**
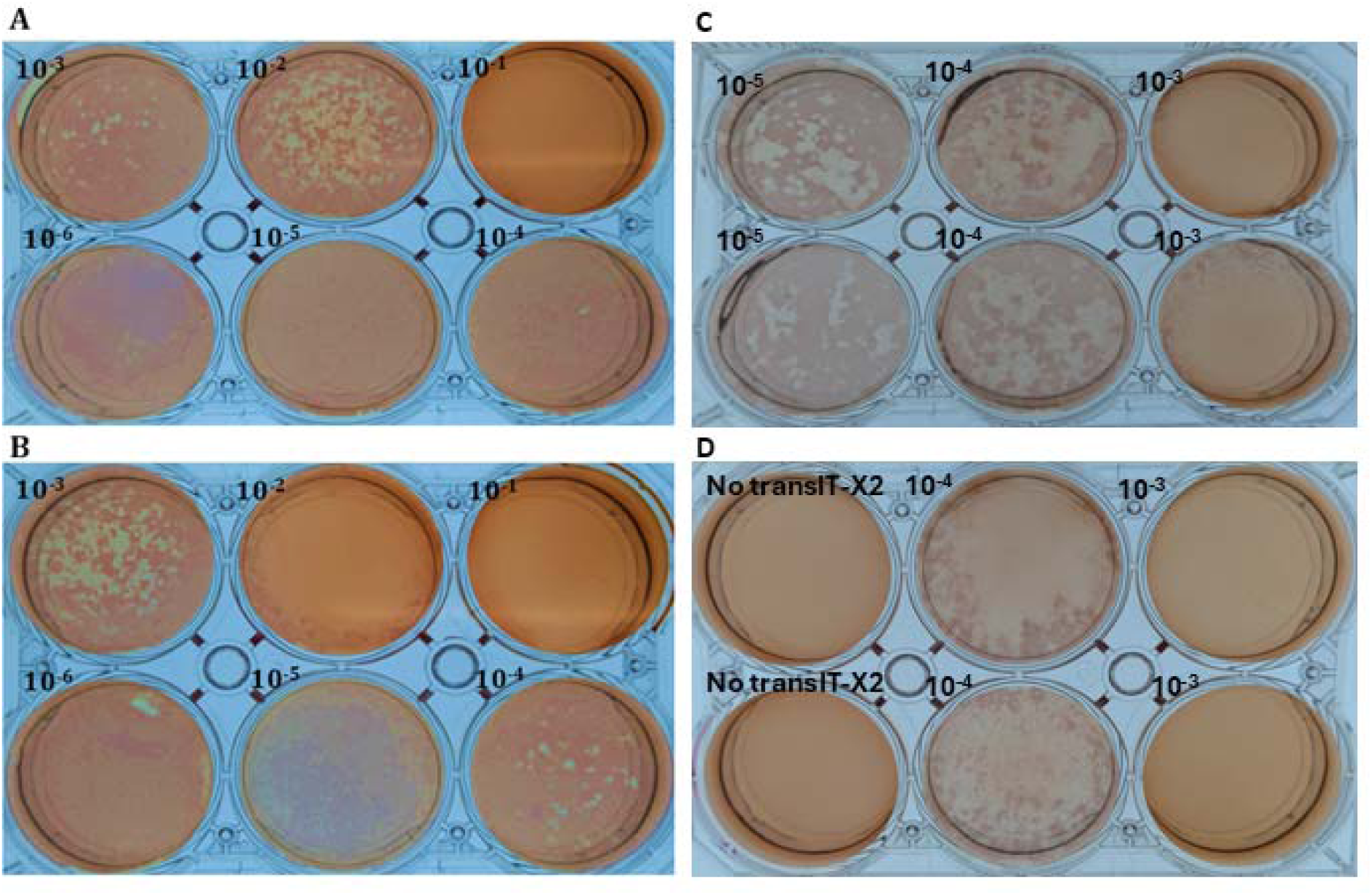
Microvesicle plaque assays demonstrating non-lytic rotavirus transmission and infectivity of MV-associated double-layered particles. Microvesicles (MVs) were isolated from MA104 cells infected with rotavirus strain SA11. MVs harvested at 8 h (A) and 22 h (B) post-infection were applied to naïve MA104 monolayers across serial dilutions, allowed to adsorb for 1 h, and overlaid with agar to assess plaque formation. Since MVs potentially contain infectious triple-layered particles, we conducted additional assays to specifically test the infectivity of MV-derived double-layered particles (DLPs). MV membranes were disrupted and the released viral particles were treated either with EDTA to remove the outer capsid of mature triple-layered particles (TLPs) (C) or with a neutralizing anti-VP7 antibody to inactivate residual TLPs (D). Treated preparations were introduced into MA104 cells by transfection using TransIT-X2^®^ reagent, and plaque formation was assessed in duplicate across serial dilutions. Control wells show parallel preparations applied in the absence of TransIT-X2^®^ reagent. At low dilutions, extensive cytopathic destruction prevented resolution of individual plaques. Quantifiable plaques were therefore observed only at higher dilutions. No plaques formed in the absence of the TransIT-X2^®^ transfection reagent, demonstrating that naked DLPs require intracellular delivery to initiate productive infection.

To confirm that the vesicular membrane is the critical element conferring infectivity to these DLPs and to demonstrate that the DLPs, not TLPs, are infectious when delivered to cells, we performed parallel control experiments in which we physically disrupted the lipid membranes of the MVs prior to inoculation, effectively stripping the lipid envelope and releasing the naked viral payload. The released viral particles were treated either with EDTA to remove the outer capsid of mature triple-layered particles (TLPs) or with a neutralizing anti-VP7 antibody to inactivate residual TLPs. Treated particles were applied to naïve MA104 monolayers across serial dilutions along with TransIT-X2 and allowed to adsorb for 1 h. The monolayers were then overlaid with agar to assess plaque formation. We observed quantifiable plaques at higher dilutions and extensive cytopathic effects at lower dilutions in both the EDTA-treated (Fig. 4C) and anti-VP7-treated (Fig. 4D) preparations (23). No plaques formed in the absence of transfection reagent. The complete loss of infectivity of treated particles following vesicle disruption confirms that the naked DLPs released from the MVs cannot independently mediate cellular entry.

Collectively, these experiments demonstrate that the host-derived lipid bilayer of the microvesicle effectively substitutes for the viral outer capsid, likely mediating cell entry through membrane fusion or endocytosis and bypassing the traditional receptor-binding requirements of free virions. Additionally, these data demonstrate that the membrane integrity of the MVs appears to be required for the non-lytic transmission and infectious capacity of rotavirus double-layered particles.

## Discussion

The traditional model of rotavirus egress holds that fully mature TLPs are released from host cells either through classical cell lysis or, in polarized cells, through a non-lytic vesicular transport mechanism (6, 7). More recent studies have shown that rotavirus can also exploit host-derived microvesicles (MVs) for cell-to-cell transmission and that rotavirus infection is associated with increased MV production (8, 9). We extend these findings by showing that the viral particles associated with released MVs are strongly enriched for immature DLPs rather than mature TLPs.

Historically, DLPs have been considered non-infectious in extracellular environments because they lack the VP4 attachment spikes and VP7 glycoprotein outer capsid required for conventional cellular entry (2). Our structural data, including TEM particle sizing and morphology assessments, differential immunogold labeling, and confocal microscopy with VP6 and VP7 protein markers, consistently demonstrate that DLPs predominate over TLPs within egressing MVs. Crucially, our plaque assays demonstrate that rotavirus-containing MVs are capable of initiating productive infections, consistent with previous observations of MV-borne rotavirus (8, 9). Our subsequent intracellular-delivery experiments further demonstrate that MV-derived DLPs are replication competent, consistent with previous studies showing that DLPs can initiate infection when delivered directly into the cytoplasm (23). While free TLPs use the VP4 attachment spike to mediate conventional cellular entry (2), MV-associated particles may instead exploit extracellular-vesicle uptake pathways. For instance, exposure of phosphatidylserine on the MV surface could facilitate entry through apoptotic mimicry and promote uptake through endocytic pathways (24, 25). Furthermore, surface-exposed phosphatidylserine can function as an immunomodulatory signal and is exploited by diverse viruses through apoptotic mimicry to dampen innate immune responses (24, 26). If phosphatidylserine is exposed on the surface of rotavirus-containing MVs, it could therefore contribute to immune evasion in addition to the passive physical protection provided by the host-derived membrane. After internalization, productive infection would depend on release of the vesicle-associated particles from the endocytic compartment, allowing transcriptionally active DLPs to reach the cytoplasm without requiring the conventional VP4/VP7-dependent entry process.

Our quantitative comparison of intracellular and MV-associated virions demonstrates that DLPs are not merely abundant within egressing MVs but are significantly enriched relative to their representation within infected cells. Whereas DLPs constituted only ∼23.4% of intracellular particles, they accounted for ∼80% of MV-associated particles, representing a striking inversion of particle-type frequency. This disparity argues against a model in which MVs simply non-selectively sample the intracellular viral population and instead suggests preferential incorporation or retention of DLPs during vesicular egress.

The mechanism underlying this enrichment remains unresolved. One possibility is that rotavirus actively directs DLPs into the MV biogenesis pathway. DLPs assembled in cytoplasmic viroplasms normally bud into the endoplasmic reticulum (ER), where they acquire the VP4 and VP7 outer capsid layers required for formation of mature TLPs (2, 3). Diversion of a subset of these particles into plasma membrane-derived vesicles before ER entry would selectively expose immature DLPs to the MV pathway and could account for their strong enrichment in extracellular vesicles. Such sorting could involve direct or indirect interactions between the exposed VP6 scaffold of DLPs and host membrane-remodeling or vesicle-sorting machinery, including components of the ESCRT pathway (10). Precedent for targeted recruitment of host vesicular machinery exists among other non-enveloped viruses. Hepatitis A virus, for example, interacts with ESCRT-associated proteins including ALIX and VPS4 to promote encapsulation within host-derived membranes (27, 28).

An alternative, although not mutually exclusive, explanation is that enrichment arises through spatial coupling between viroplasms and sites of MV biogenesis. If MV-forming membrane domains preferentially intersect with viroplasms or with trafficking routes carrying newly assembled DLPs toward the ER, DLPs could be selectively captured before outer-capsid maturation. High-resolution spatiotemporal imaging will therefore be important for distinguishing molecular sorting from spatially driven enrichment. Co-localization of MV-associated markers such as flotillin-2 with the viroplasm proteins NSP2 and NSP5, together with ER markers, could reveal whether sites of vesicle formation are physically associated with DLP production or trafficking. Perturbation of classical ER-directed maturation, for example through manipulation of NSP4 function, could further determine whether altered access to the ER changes the proportion of DLPs entering the MV-mediated egress pathway.

The robust increase in the lipid raft-associated protein flotillin-2, a structural component associated with MV formation, further suggests that rotavirus infection substantially alters host membrane dynamics (Fig. 1). Together with the preferential enrichment of DLPs within MVs, this observation raises the possibility that rotavirus does more than passively exploit basal vesicle shedding and may instead reprogram host membrane-remodeling pathways to facilitate non-lytic viral dissemination (29, 30). Increased MV production could provide a continuously available pool of host-derived membranes capable of packaging infectious DLPs.

However, increased vesiculation may also represent a host response to infection rather than a virus-directed process. Infected cells can enhance extracellular vesicle release to export damage-associated molecular patterns, viral antigens, and innate immune signaling molecules that alert neighboring cells and promote antiviral defenses (28, 31). Under this model, rotavirus may exploit an infection-induced host vesiculation response while simultaneously biasing the viral cargo toward DLPs. The strong enrichment of DLPs within MVs therefore supports a model in which vesicular rotavirus egress is a selective process, although determining whether that selectivity results from direct molecular sorting, spatial organization of viral replication and membrane-remodeling pathways, or a combination of both will require further investigation.

If rotavirus actively drives the preferential packaging of DLPs into MVs, it may represent an evolutionary adaptation for efficient host exploitation. We speculate that circumventing the energy-intensive assembly of the outer capsid could allow the virus to redirect the host’s metabolic resources toward genome replication and inner core synthesis. By minimizing the energetic investment required per genome, the virus maximizes the total yield of infectious progeny generated by a single cell. Crucially, this metabolic economy fundamentally alters the stoichiometry of the infection event. By bypassing the protein investment normally required to construct an individual outer capsid for every genome, the virus can instead package multiple DLPs, frequently exceeding five particles per vesicle, into a single host-derived envelope (8).

Consequently, a single vesicular entry event guarantees a high localized multiplicity of infection (8). Crucially, this dense packaging mechanism observed *in vitro* directly drives the massive, localized inocula required to overcome early cellular replication barriers and enhance disease severity during *in vivo* fecal-oral transmission (8). This simultaneous delivery of multiple genomes may also facilitate the complementation of deleterious mutations through shared replication machinery and establish an intracellular environment highly conducive to genetic reassortment and accelerated viral evolution (8, 15, 32–35). Furthermore, cloaking rotavirus within a host-derived membrane shields virions from neutralizing antibodies targeted against outer capsid proteins and may protect the viral payload from luminal proteases and harsh environmental stressors, such as chlorine disinfection, thereby enhancing environmental stability and transmission rates (9, 11, 36).

Given the potential advantages of collective transmission, why do rotaviruses continue to produce fully mature TLPs? This behavior is especially puzzling given evidence that vesicle-cloaked rotavirus clusters are highly effective units for inter-organismal fecal-oral transmission and remain intact during passage through the gastrointestinal tract (8). By contrast, free TLPs shed in stool are significantly less infectious *in vivo*, likely because of transit dilution and increased exposure to intestinal proteases or mucosal antibodies (8). One possibility is that maintaining an independent free-virus pathway periodically imposes low-multiplicity transmission bottlenecks that limit the persistence of defective or interfering viral genotypes. This possibility has direct precedent in rotavirus. Serial passage of bovine rotavirus at high multiplicity selected viruses carrying rearranged genome segments, whereas standard viruses predominated during low-multiplicity passage (37). Similarly, serial high-multiplicity passage of porcine rotavirus produced a defective and interfering population carrying a rearranged NSP2-NSP5 genome segment that depended on coinfection with replication-competent helper virus and interfered with parental-virus replication; this defective population rapidly disappeared following low-multiplicity passage (38).

Thus, the high cellular multiplicity generated by vesicular delivery may provide conditions that permit defective or interfering genomes to persist through complementation with functional viral genomes. By contrast, transmission as individual TLPs may periodically impose a genetic bottleneck that favors genomes capable of completing the viral life cycle independently and reduces the persistence of variants dependent on coinfection. In this model, vesicle-mediated and free-virus transmission represent complementary strategies: collective transmission provides immediate benefits through the simultaneous delivery of multiple genomes, whereas individual TLP transmission may help preserve viral population integrity over longer evolutionary timescales. More generally, alternating between high- and low-multiplicity transmission could balance the short-term advantages of collective infection against the genetic costs associated with sustained high multiplicity, including the accumulation of defective or interfering genomes (39–42). Whether maintenance of the free-TLP transmission pathway evolved specifically in response to these costs remains unknown.

This revised understanding of non-lytic, MV-mediated egress expands our view of rotavirus transmission and may have implications for vaccination and treatment. Current live oral rotavirus vaccines elicit multifaceted immune responses, including neutralizing antibodies directed against the outer capsid proteins VP4 and VP7 (43). Because vesicular cloaking can reduce the accessibility of viral capsid proteins to neutralizing antibodies, MV-associated transmission could provide rotavirus with a mechanism of partial immune evasion (8, 9). Additionally, although rotavirus was historically considered a strictly enteric pathogen, it is now well established that infection is frequently accompanied by antigenemia and viremia and can involve replication in extra-intestinal tissues (12–14). The mechanisms that enable this systemic dissemination remain incompletely understood. Free virions circulating outside the gastrointestinal tract remain directly exposed to rotavirus-specific serum antibodies and other systemic immune effectors, which contribute to the control and clearance of rotavirus antigenemia and viremia (44).

Our findings raise the possibility that host-derived MVs could contribute to extra-intestinal dissemination by providing a membrane barrier around replication-competent DLPs. Such cloaking could reduce direct antibody access to viral antigens, including the highly immunogenic VP6 scaffold (9, 45). Vesicle-associated DLPs could therefore be partially protected from circulating antibodies during dissemination through lymphatic or circulatory pathways (9), potentially facilitating access to extra-intestinal sites such as the liver and central nervous system (12, 14). Although the role of MV-mediated transport in systemic infection and vaccine escape remains to be established *in vivo*, these findings, together with previous evidence for vesicle-associated rotavirus transmission and protection from neutralizing antibodies, suggest that vesicle-associated transmission should be considered when investigating mechanisms of rotavirus dissemination, immune evasion, and potential therapeutic intervention (8, 9, 33).

## Materials and Methods

### Cell Culture and Viral Propagation

Human colon carcinoma (HT-29) and African green monkey kidney (MA104) cell lines were obtained from the American Type Culture Collection (ATCC). HT-29 cells were passaged every five days in McCoy’s 5A medium (Cytiva), while MA104 cells were passaged every three days in Medium 199 (Cytiva). Both media were supplemented with 10% fetal bovine serum (FBS; Cytiva) and 1% penicillin-streptomycin-amphotericin B (Sigma Aldrich). Cells were maintained at 37°C in a humidified incubator with 5% CO_2_. To infect HT-29 and MA104 cells, Rhesus rotavirus (RRV; gifted by Harry Greenberg) and simian rotavirus (SA11; gifted by Susan Estes) strains were used, respectively. Rotaviruses were activated with trypsin at a final concentration of 10 µg/mL for 1 h at 37°C.

### Vesicle Isolation

HT-29 and MA104 monolayers grown in T75 flasks were washed twice with 1x PBS and infected with RRV and SA11 or RRV, respectively, at an MOI of 1. Adsorption was carried out for 1 h at 37°C in 5% CO_2_. Following adsorption, monolayers were washed twice with 1x PBS and maintained in serum-free McCoy’s 5A (for HT-29) or Medium 199 (for MA104). Vesicles were collected at different points post-infection. The supernatant from infected cells was collected and centrifuged at 2,500 x g for 10 min to remove large cell debris. The supernatant was then processed using a tangential flow filtration (TFF) system equipped with an Omega^™^ 300K membrane (Minimate^™^ TFF capsule, Pall Corporation) to retain particles within the desired size range, including microvesicles approximately 100–1,000 nm in diameter. The retentate was collected and centrifuged at 17,000 × g for 35 min to pellet the microvesicles. The pellet was washed once with cold 1x PBS by centrifugation at the same speed for 30 min. After removal of the supernatant, the final pellet was resuspended in 100–200 µL of cold 1x PBS and stored at 4°C for future experiments.

### Western Blotting

MV isolation and total production were evaluated by Western blotting for flotillin-2, an established MV marker (16). MA104 cells were infected with SA11 rotavirus at an MOI of 1. Microvesicles were isolated after 4, 6, 8, 12, 16, and 24 h post-infection. Total protein concentration was quantified using a Qubit Protein Broad Range Assay Kit (Thermo Fisher Scientific). Samples were resuspended in 4x Laemmli sample buffer (Bio-Rad) and resolved on a 4–20% SDS-PAGE gel under non-reducing conditions. Proteins were transferred to a nitrocellulose membrane and, to validate and quantify MV production, probed with an anti-flotillin-2 primary antibody followed by a near-infrared (NIR) anti-rabbit secondary antibody. Blots were visualized using an Odyssey DLx Imaging System (LI-COR Biosciences), and total protein normalization was performed using Revert 700 Total Protein Stain (LI-COR).

### Transmission Electron Microscopy

Four microliters of vesicles isolated at 20 h post-infection from SA11-infected MA104 cells or from RRV-infected HT-29 cells were adsorbed onto carbon-coated 200-mesh copper grids (Ted Pella) for 1 min at room temperature (RT). Excess liquid was soaked away with a small piece of blotting paper. They were then negatively stained by the single-drop method with 4 µL of a 2% solution of uranyl acetate dissolved in deionized water and incubated at RT for 30 sec. Several 10–15 µL water droplets were placed on a separate piece of Parafilm, and grids were positioned sample-side down on each droplet to sequentially wash away background staining three times. The grids were then air dried for at least 15 minutes before they were imaged. The imaging was performed using a JEOL JEM-1400Flash transmission electron microscope operating at 100 kV, and micrographs were captured at 20,000-50,000x magnification using an AMT sCMOS 16Mb digital camera.

### Microvesicle-Derived Rotavirus Plaque Assays

The infectivity of rotavirus associated with microvesicles (MVs) was assessed by plaque assay. To neutralize residual free triple-layered particles (TLPs), MVs isolated from SA11-infected MA104 cells were incubated at 37°C for 90 minutes with 6 µg/mL of a neutralizing anti-VP7 outer capsid glycoprotein antibody (Anti-Rotavirus A/RV-A Outer Capsid Glycoprotein VP7, Cat. No. VVV20001, Antibody System) (19). The treated MV preparation was applied to confluent MA104 cell monolayers in 6-well plates and allowed to adsorb for 1 h at 37°C in 5% CO₂, with plates manually rocked every 15 min. Following adsorption, monolayers were overlaid with a 1:1 mixture of 1.4% SeaPlaque^®^ agarose (Lonza) and 2× EMEM (Quality Biologicals). Plates were incubated at 37°C in 5% CO₂ until plaques developed and were subsequently stained with neutral red. As a control for the requirement of an intact MV membrane, an aliquot of the MV preparation was disrupted using RIPA buffer and sonication. RIPA buffer was subsequently removed using HiPPR^™^ Detergent Removal Spin Columns (Thermo Scientific) prior to application of the preparation to MA104 cell monolayers. Each experimental condition was evaluated in duplicates across three independent experiments.

To evaluate the replication competence of DLPs recovered from MVs, MVs isolated from SA11-infected MA104 cells were disrupted by adding Vertrel™ XF at a 1:1 ratio. The mixture was vortexed at high speed for 3 minutes using a Fisher Scientific vortex mixer and immediately placed on ice for 3 minutes. This extraction cycle was repeated three times to fully release the enclosed viral particles, which were subsequently collected from the upper aqueous phase. The recovered viral suspension was divided into two aliquots and subjected to independent treatments designed to eliminate infectivity attributable to mature TLPs. The first aliquot was treated with EDTA to a final concentration of 10 mM and incubated at 37°C for 30 minutes to remove the VP4/VP7 outer capsid from mature TLPs by chelating the calcium ions required for outer-capsid stability (19, 46). Following treatment, EDTA was quenched by addition of CaCl₂ to a final concentration of 13 mM. The second aliquot was incubated with neutralizing anti-VP7 antibody (Anti-Rotavirus A/RV-A Outer Capsid Glycoprotein VP7, Cat. No. VVV20001, Antibody System) to neutralize naturally occurring TLPs remaining in the MV-derived viral preparation. Removal of the VP7 outer capsid following EDTA treatment was evaluated by Western blotting for the rotavirus structural proteins VP6 and VP7.

Because free DLPs lack the VP4/VP7 outer capsid required for conventional cellular entry, the EDTA-treated and anti-VP7-treated preparations were introduced into confluent MA104 cell monolayers by lipid-mediated transfection. Intracellular delivery was performed as previously described for evaluating the infectivity of transfected rotavirus DLPs (1), utilizing the TransIT-X2® Dynamic Delivery System (Mirus Bio) as the transfection reagent at a 1:10 (v/v) reagent-to-sample ratio per well. Parallel preparations were applied to MA104 monolayers in the absence of TransIT-X2 as negative controls for spontaneous entry of the released DLPs. At 24 h post-transfection, culture medium was aspirated and the monolayers were overlaid with 1.4% SeaPlaque™ agarose. Plaques were visualized and quantified by neutral red staining on day 6 post-transfection from two independent replicates at dilutions permitting resolution of individual plaques. While alternative protocols have utilized Lipofectamine 2000, the present study employed the less cytotoxic TransIT-X2 reagent. Furthermore, this experiment investigated whether inactivated virus particles can successfully establish an infection when delivered intracellularly via extracellular vesicles.

### Immunoelectron Microscopy Staining Using Gold-Conjugated Secondary Antibodies

Immunoelectron microscopy was used to visualize microvesicle–associated rotavirus particles. MVs isolated from SA11-infected MA104 cells were adsorbed onto carbon-coated 200-mesh copper grids (Ted Pella) for 5 minutes and subsequently treated with Intercept^®^ blocking buffer (LICORbio) for 30 minutes to reduce nonspecific binding. The grids were then incubated with primary antibodies against VP6 (a marker for double-layered particles, DLPs) and VP7 (a marker for triple-layered particles, TLPs) for 2 hours. DLP antibodies were mouse monoclonal antibodies against rotavirus inner capsid protein VP6 (clone A2; Novus Biologicals, Cat#NB110-37243, 1:1,000 dilution in blocking buffer), and TLP antibodies were rabbit polyclonal antibodies against rotavirus outer capsid glycoprotein VP7 (Cusabio, Cat# CSB-PA420195ZA01RIU, 1:1,000 dilution in blocking buffer). The grids were washed 3 times for 3 minutes in a droplet of wash buffer (100 mL 1× PBS, 100µL Tween 20, 100µL 30% bovine serum albumin (BSA), IgG-free). Washing was followed by incubation with 8-nm (Cytodiagnostics; Goat anti-Mouse IgG (H+L), Cat#AC-10-02-05), and 6-nm (Invitrogen; Goat anti-Rabbit IgG (H+L), Cat#A-31565) gold-conjugated secondary antibodies for VP6 and VP7, respectively, for 1 hour. The grids were washed another 3 times for 3 minutes in a droplet of wash buffer, followed by negative staining with a 2% uranyl acetate solution for 5 minutes. The grids were washed once in a droplet of water for 3 seconds and wicked dry. A JEOL 1400Flash TEM was used to perform immunoelectron microscopy at 100 kV with an AMT sCMOS 16Mb digital camera.

### Immunofluorescence Labeling and Cell Imaging

We seeded MA104 cells into 35-mm glass-bottom dishes (10-mm well, Cellvis) and incubated them for 24 h at 37°C with 5% CO₂. After infection with SA11 virus stock, we fixed the monolayers at specified time points using 4% PFA in 1x PBS for 10 min at room temperature, followed by three washes in 1x PBS. We incubated the fixed samples with primary antibodies in incubation buffer (1x PBS, 1% BSA, and 0.2% saponin) for 1 h at RT or overnight at 4°C. After washing the cells four times with 1x PBS, we applied fluorescently tagged secondary antibodies diluted in the same buffer and incubated them for 1 h at RT in the dark. We then washed the samples three times with 1x PBS and stained the cells with Cytoliner^TM^ 410/450, a fixed cell membrane stain per the manufacturer’s protocol (Biotium). All samples were imaged using a Zeiss LSM900 confocal laser scanning microscope.

## Data Availability

All data supporting the findings of this study are available within the article and its Supporting Information. Underlying data, including uncropped Western blots, transmission electron microscopy images, particle-size measurements and classifications, immunogold and confocal microscopy images, and plaque-assay data, are available from the corresponding author upon reasonable request.

## Acknowledgments

The authors are grateful to Nihal Altan-Bonnet and John Patton for their insightful discussions and valuable feedback on this work. Rhesus rotavirus and simian rotavirus SA11 were gifts from Harry Greenberg and Susan Estes, respectively. This work was funded in part by grants from the NSF Division of Environmental Biology (Award #2517851) and the Ira Spar Biosciences Laboratory at Queens College.

## Author Contributions

- designed research (ZMI, JJD)
- performed research (ZMI, JL, AC, AS, EC, ISM, KN)
- analyzed data (ZMI, JL, AC)
- wrote the paper (ZMI, JL, JJD)

## Classification

Microbiology

