## Supporting Information for "Selective Packaging of Rotavirus Double-Layered Particles into Host Microvesicles"

John J. Dennehy

**This PDF file includes:**

Figure S1

Legend for Movie S1

SI Reference

**Other supporting materials for this manuscript include the following:**

Movie S1

**Figure S1**

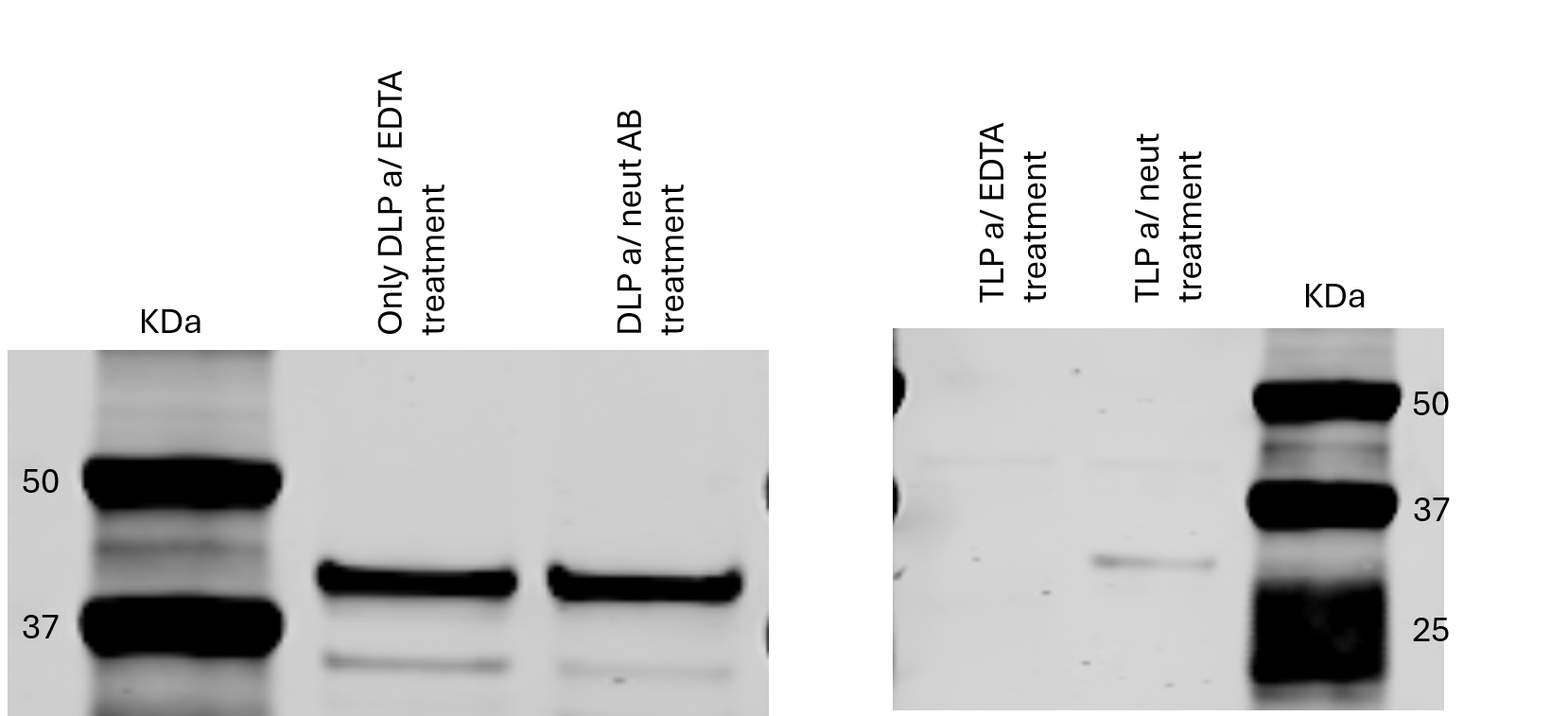

(A)

(B)

Fig. S1. Characterization of treated rotavirus particle fractions by Western blot analysis. Immunoblotting was performed on EDTA-treated and neutralizing antibody-treated viral samples isolated from microvesicles to assess outer-capsid integrity and structural composition. (A) Both experimental groups exhibit distinct, robust bands corresponding to the inner capsid protein VP6 (~40–45 kDa), confirming the uniform presence and integrity of double-layered particles (DLPs) across treatments. (B) Analysis of the outer capsid glycoprotein VP7 (~34–37 kDa) showing that the EDTA-treated sample demonstrates a complete absence of VP7 signal, validating successful calcium chelation and triple-layered particle (TLP) stripping. Conversely, the sample treated with anti-VP7 neutralizing antibody reveals trace immunoreactivity within the 34–37 kDa range, indicating a residual population of intact or antibody-complexed TLPs.

Movie S1. Confocal imaging movie of viral budding. MV-associated particles were analyzed via dual-color immunofluorescence at 14h post-infection. TLPs (anti-VP7) were visualized with Alexa Fluor 647 (red); DLPs (anti-VP6) were visualized with Alexa Fluor 488 (green). Cytoliner^TM^ 410/450, fixed cell membrane stain (Biotium) was used to stain the cells. Confocal analysis reveals a prominent abundance of green DLP signal relative to red TLP signal at peripheral budding sites.
